# Functional divergence of DSCAM family in vertebrates through domain-specific evolutionary pressures

**DOI:** 10.64898/2026.04.14.718097

**Authors:** Koichi Hashizume, Yusuke Watanabe, Hiroki Oota, Mikio Hoshino

## Abstract

The Down syndrome cell adhesion molecule (DSCAM) family, conserved across metazoans, plays key roles in neural development by mediating cell-cell recognition. Invertebrate Dscam evolved extensive molecular diversity through isoform diversification, whereas the vertebrate paralogs DSCAM and DSCAML1 followed a distinct evolutionary trajectory. However, how these vertebrate paralogs evolved after duplication, particularly with respect to functional divergence, remains poorly understood. Here, we investigated the evolutionary history and post-duplication divergence of these paralogs using phylogenetic, molecular evolutionary, sequence comparison, and transcriptomic analyses. Our phylogenetic analyses suggest an ancestral gene duplication predating the split between gnathostomes and cyclostomes. We found distinct patterns of selective constraint between the paralogs, particularly in the intracellular domain. In tetrapods, the intracellular domain of DSCAM showed strengthened purifying selection, whereas no comparable reinforcement was evident for DSCAML1, despite strong constraint in mammals. We also found distinct patterns of lineage- and site-specific positive selection between DSCAM and DSCAML1. Consistent with these evolutionary differences, comparative analysis of the intracellular domains revealed distinct repertoires of short linear motifs (SLiMs) predicted to mediate protein-protein interactions. Reanalysis of published transcriptomic data further suggested distinct downstream responses elicited by the intracellular domains of DSCAM and DSCAML1. Together, these findings suggest that post-duplication functional divergence of vertebrate DSCAM paralogs may have contributed to the evolution of molecular mechanisms underlying vertebrate neural development and circuit formation.

## Introduction

Cell adhesion molecules (CAMs) are essential components of the molecular systems that couple selective cell-surface recognition to appropriate intracellular responses during neural development. In the nervous system, CAM-mediated interactions regulate neuronal migration, neurite outgrowth and guidance, synaptic target selection, and synaptogenesis, thereby contributing to the establishment of functional neural circuits (Moreland and Poulain, 2022). The repertoire and functional diversity of CAMs have expanded throughout metazoan evolution, concomitant with the increasing complexity of nervous systems (Cortés et al., 2023). This evolutionary diversification has facilitated refined cell-cell recognition and synaptic specificity, likely contributing to the emergence of sophisticated neural networks.

The *Down syndrome cell adhesion molecule* (*Dscam*) gene encodes a single-pass transmembrane molecule belonging to the immunoglobulin superfamily. Its domain architecture is evolutionarily conserved from mammals to the fruit fly *Drosophila melanogaster*, consisting of an N-terminal extracellular region with ten immunoglobulin-like domains and six fibronectin type III domains, followed by a transmembrane region and a C-terminal intracellular domain (Yamakawa et al., 1998). In *Drosophila*, four paralogous *Dscam* genes (*Dscam1-4*) have been identified, among which *Dscam1* is particularly notable for its extensive isoform diversity generated through alternative splicing (Schmucker et al., 2000; Neves et al., 2004). *Dscam1* produces 12, 48, and 33 alternative isoforms for its second, third, and seventh Ig domains, respectively, resulting in 19,008 potential extracellular domain variants (Schmucker et al., 2000). These isoforms are stochastically expressed in individual neurons and mediate isoform-specific homophilic interactions, providing a molecular code that distinguishes ‘self’ from ‘non-self’. This mechanism enables neuronal self-avoidance and proper neural circuit assembly (Wojtowicz et al., 2004, 2007; Zhan et al., 2004; Hattori et al., 2007; Hughes et al., 2007; Matthews et al., 2007; Miura et al., 2013). The extensive isoform diversity of *Dscam* is evolutionarily conserved across arthropods, although the mechanisms generating this diversity vary among lineages (Yue et al., 2016; Hou et al., 2022).

In vertebrates, the *Dscam* family comprises two paralogous genes, *Dscam* and *Dscaml1* (Yamakawa et al., 1998; Agarwala et al., 2001). Unlike their arthropod counterparts, vertebrate *Dscam* and *Dscaml1* do not generate extensive isoform diversity. Instead, isoform diversity in vertebrate neural systems is mediated primarily by the clustered protocadherins (cPcdhs), which contribute to neuronal self-recognition and self-avoidance (Lefebvre et al., 2012; Thu et al., 2014; Kostadinov and Sanes, 2015; Chen et al., 2017; Mountoufaris et al., 2017). Nevertheless, DSCAM and DSCAML1 are functionally important in vertebrate neural development (Hizawa et al., 2024). Similar to *Drosophila* Dscam, vertebrate DSCAM and DSCAML1 mediate isoneuronal and homotypic self-avoidance and regulate neurite arborization and mosaic tiling (Fuerst et al., 2008, 2009; Yamagata and Sanes, 2008; Garrett et al., 2018). Recent studies have further revealed diverse roles for these proteins in vertebrate neural development. In the developing mouse cerebral cortex, both genes influence radial neuronal migration (Zhang et al., 2015; Mitsogiannis et al., 2020; Yang et al., 2022). We also reported that DSCAM facilitates the delamination of neuronal progenitor cells from the ventricular surface in the midbrain (Arimura et al., 2020), and promotes perisynaptic localization of the glutamate transporter GLAST in cerebellar Purkinje cells, thereby ensuring efficient glutamate clearance (Dewa et al., 2024). Moreover, DSCAM and DSCAML1 have been shown to promote cell death in mouse retina and zebrafish hypothalamus, respectively (Li et al., 2015; Ma et al., 2023). Beyond these roles, both proteins have also been implicated in transcriptional regulation: following proteolytic cleavage by γ -secretase, their intracellular domains translocate to the nucleus and modulate the expression of genes involved in neuronal differentiation, synaptogenesis, and synaptic function (Sachse et al., 2019). Collectively, these findings suggest that vertebrate DSCAM family proteins may have evolved to fulfill multiple roles that support neural circuit development and organization in a manner distinct from that of arthropods.

Gene duplication is widely recognized as a key mechanism driving the emergence of novel gene functions during evolution (Ohno, 1970). Recent studies suggest that DSCAM and DSCAML1 are not fully functionally equivalent in mammals. For instance, overexpression of these genes differentially affects synapse development in primary hippocampal neurons and migration of cortical interneurons (Sachse et al., 2019; Mitsogiannis et al., 2020). These findings raise the possibility that the two vertebrate paralogs were shaped by distinct patterns of molecular evolution after duplication. However, the evolutionary processes underlying these apparent functional differences remain poorly understood across vertebrates, partly because previous studies have focused primarily on arthropod *Dscam* diversification or vertebrate datasets with limited taxonomic sampling (Crayton et al., 2006; Brites et al., 2008; Lee et al., 2010; Cao et al., 2018; Wiseglass and Rubinstein, 2024).

Here, we investigated the evolutionary history of DSCAM and DSCAML1 across vertebrates using broad phylogenetic sampling, focusing on domain-specific patterns of molecular evolution. We reconstructed the phylogenetic relationships of these paralogs, compared the selective pressures acting on their extracellular and intracellular domains, analyzed intracellular-domain sequence variation, and reanalyzed published transcriptomic data. Our results indicate that DSCAM and DSCAML1 followed asymmetric patterns of molecular evolution and further suggest functional divergence, particularly in their intracellular domains. Together, these findings suggest that such asymmetric post-duplication evolution may have contributed to the evolution of molecular mechanisms underlying vertebrate neural development and circuit formation.

## Methods

### Sequence data collection

Coding sequences (CDS) for the *Dscam* and *Dscaml1* genes were obtained from Ensembl v.110 and NCBI RefSeq release 221, covering a broad range of representative vertebrate taxa. The dataset comprised sequences from 78 species: 39 mammals, 12 birds, 8 reptiles, 4 amphibians, 9 ray-finned fishes, 2 cartilaginous fishes, 2 cyclostomes, as well as an echinoderm and an arthropod (Supplementary Table S1). To verify orthologous relationship for *Dscam* family members in cyclostomes, the echinoderm, and the arthropod, we used OrthoFinder v.3.0 (Emms and Kelly, 2019). In ray-finned fishes, gene duplication events have led to the presence of two distinct *Dscam* paralogs, termed *dscama* and *dscamb*, while no duplication has been identified for *Dscaml1*. Both *Dscam* paralogs in teleosts were retained for downstream analyses.

### Molecular phylogenetic analyses and synteny analysis

Nucleotide sequences of *Dscam* and *Dscaml1* genes were translated into amino acid sequences and aligned using the MUSCLE algorithm implemented in MEGA X (Kumar et al., 2018). To improve the quality of phylogenetic inference, poorly aligned regions and gap-rich sites were trimmed using trimAl v.1.4.rev22 (Capella-Gutiérrez et al., 2009) with the gappyout option. Maximum likelihood (ML) phylogenetic trees were constructed with IQ-TREE2 (Minh et al., 2020) using the best-fitting substitution model determined by ModelFinder (Kalyaanamoorthy et al., 2017). Bayesian inference trees were generated using MrBayes v.3.2.7 (Ronquist et al., 2012), running four simultaneous chains for 1,000,000 generations with sampling every 500 generations. The first 25% of sampled trees were discarded as burn-in, and convergence was considered achieved when the average standard deviation of split frequencies dropped below 0.01.

The genomic context of *Dscam* family genes was analyzed by examining the arrangement of protein-coding genes flanking *Dscam* and *Dscaml1* loci in selected representative vertebrate species: human (*Homo sapiens*), chicken (*Gallus gallus*), zebrafish (*Danio rerio*), thorny skate (*Amblyraja radiata*), sea lamprey (*Petromyzon marinus*), and hagfish (*Eptatretus burgeri*). Genomic data for these species were accessed via the Ensembl Genome Browser and the NCBI Genome Data Viewer. The genomic regions surrounding the *Dscam* and *Dscaml1* loci were manually inspected to identify neighboring genes.

To assess amino acid conservation within the DSCAM family, the MUSCLE-aligned sequences were analyzed in JalView (Waterhouse et al., 2009), and conservation scores for each alignment position were calculated using the AMAS method (Livingstone and Barton, 1993).

### Molecular evolutionary analyses

Transmembrane regions were predicted from the translated amino acid sequences using the TMbed protein language model (Bernhofer and Rost, 2022). Extracellular domains were defined as regions N-terminal to the transmembrane segment, and the intracellular domains as regions C-terminal to it.

Pairwise nonsynonymous-to-synonymous substitution rate ratios (dN/dS) were estimated for each domain of DSCAM and DSCAML1 using the modified Nei–Gojobori (proportional) method implemented in MEGA X. Alignment sites containing gaps were excluded from the analysis to avoid artifacts. Statistical significance was assessed using the Kruskal–Wallis rank-sum test, followed by post hoc Dunn’s tests with Bonferroni correction for multiple comparisons.

We used RELAX (Wertheim et al., 2015) implemented in DataMonkey webserver (Weaver et al., 2018) to test whether the overall strength of selection was relaxed or intensified along a predefined set of branches. The tetrapod stem branch and all descendant branches excluding mammals were designated as the test set, whereas ray-finned fishes together with cartilaginous fishes were used as the reference set. Multiple sequence alignments for each domain were trimmed using trimAl with the gappyout option to eliminate poorly aligned regions. RELAX estimates a selection intensity parameter (k), where k < 1 indicates relaxation and k > 1 indicates intensification of selection on the test branches relative to the reference branches by scaling the distribution of ω values. Statistical support was assessed using a likelihood ratio test (LRT) comparing a null model with k fixed at 1 to an alternative model in which k was estimated from the data.

To detect signals of positive selection acting on specific lineages, we employed the branch-site model (Model A vs. Model A null) implemented in CodeML from the PAML package v4.9 (Yang, 2007). Multiple sequence alignments for each domain were trimmed using trimAl with the gappyout option to eliminate poorly aligned regions. For each gene, ML phylogenetic trees based on the extracellular domain sequences were used to represent species relationships. Foreground branches were designated for key evolutionary nodes, including the most recent common ancestors of osteichthyans, tetrapods, amniotes, and mammals. LRTs were performed to compare Model A and its null counterpart, and significance was assessed based on the χ² distribution with one degree of freedom.

### *In silico* protein analysis

To assess functional divergence among domains of the DSCAM family in tetrapods, we employed DIVERGE v3.0 (Gu et al., 2013). Multiple protein sequence alignments of the extracellular and intracellular domains were constructed using the MUSCLE algorithm implemented in MEGA X, including sequences from one amphibian (*Xenopus tropicalis*), two reptiles (*Hemicordylus capensis*, *Pseudonaja textilis*), two birds (*Gallus gallus*, *Serinus canaria*), and seven mammals (*Balaenoptera musculus*, *Bos taurus*, *Canis lupus familiaris*, *Homo sapiens*, *Macaca mulatta*, *Monodelphis domestica*, *Mus musculus*). The coefficients of functional divergence, θ of Type I and Type II, were calculated as measures of functional differentiation. The value of θ ranges from 0 (no functional divergence) to 1 (complete divergence). A significant deviation of θ from 0 indicates significant functional divergence between gene clusters. Type I functional divergence refers to functional differentiation caused by a shift in evolutionary rate among sites, whereas Type II functional divergence refers to cases where the evolutionary rate remains similar but amino acid properties have changed, leading to altered functional constraints. Type-I functional divergence was estimated using the model-free method and Type-II functional divergence was calculated using the simplified ML method. Statistical significance was evaluated by calculating the Z-score (θ/SE) under the assumption of a normal distribution.

Predicted full-length protein structures for human DSCAM and DSCAML1 were obtained from the AlphaFold Protein Structure Database (Varadi et al., 2024). The intracellular domain structures were further predicted using ColabFold v1.5.5 (Mirdita et al., 2022). To assess intrinsic disorder, we performed disorder prediction analyses using IUPred3 (Erdős et al., 2021) with full-length amino acid sequences of human DSCAM and DSCAML1.

Short linear motifs (SLiMs) were identified using the Eukaryotic Linear Motif (ELM) database (Kumar et al., 2024). SLiMs in the ELM database are categorized into six functional classes: cleavage sites (CLV), degradation sites (DEG), docking sites (DOC), ligand-binding sites (LIG), post-translational modification sites (MOD), and subcellular targeting motifs (TRG). We focused on SLiMs located within the intracellular domains of human DSCAM and DSCAML1 that belong to the DOC and LIG classes. Identified SLiMs were classified into two groups: (1) conserved SLiMs, which are present at the same aligned positions in both DSCAM and DSCAML1 and are inferred to have arisen prior to the gene duplication event, and (2) paralog-specific SLiMs, which are found in only one of the two paralogs. To examine the evolutionary conservation of these motifs, intracellular domain SLiMs were also identified for DSCAM and DSCAML1 in mouse (*Mus musculus*), zebrafish (*Danio rerio*), and thorny skate (*Amblyraja radiata*). Human paralog-specific SLiMs were then compared with those identified in these species, and further classified into four categories based on their conservation patterns: (1) gnathostome-conserved paralog-specific SLiMs (G-PS), present in all four species, (2) osteichthyan-conserved paralog-specific SLiMs (O-PS), shared among human, mouse and zebrafish but absent in thorny skate, (3) mammal-conserved paralog-specific SLiMs (M-PS), found only in human and mouse, and (4) non-conserved paralog-specific SLiMs (non-conserved PS), defined as all remaining paralog-specific SLiMs that do not meet the criteria for categories (1)–(3).

### Bulk RNA-seq data analysis

Bulk RNA-seq data from HEK293 t-Rex-Flp-In cell lines stably expressing YFP-tagged intracellular domains of DSCAM and DSCAML1, or expressing cytoplasmic or nuclear YFP alone as controls, were retrieved from the Gene Expression Omnibus (GEO) under accession number GSE122568 (Sachse et al., 2019). Differentially expressed genes (DEGs) between DSCAM vs. nuclear YFP and DSCAML1 vs. nuclear YFP were identified using DESeq2 v.1.42.1 (Love et al., 2014), applying a threshold of fold change > 2 and adjusted p-value ≤ 0.05. To minimize false positives, DEGs identified between cytoplasmic YFP and nuclear YFP were excluded from further analysis, as described previously (Sachse et al., 2019).

To define paralog-specific DEGs, we first identified genes that were significantly upregulated or downregulated in cells expressing DSCAM or DSCAML1, respectively. Upregulated genes from each condition were then compared to identify DSCAM- and DSCAML1-specific upregulated genes; the same procedure was applied to downregulated genes. Finally, paralog-specific DEGs were defined as the union of paralog-specific upregulated and downregulated genes. Gene Ontology (GO) enrichment analysis for Biological Process terms was conducted using ShinyGO v.0.82 (Ge et al., 2020) and PANTHER v.19.0 (Thomas et al., 2022), with a false discovery rate (FDR) p-value threshold of 0.05.

## Results

### Phylogeny of the DSCAM family in vertebrates

We first investigated the phylogenetic relationships within the vertebrate DSCAM family and its close homologs. A total of 160 sequences were collected from 78 species (Supplementary Table S1), including cyclostomes (sea lamprey and hagfish), an echinoderm (sea urchin), and an arthropod (fruit fly). No orthologs were identified in the cephalochordate amphioxus or the urochordate sea squirt. Using the maximum likelihood (ML) method, we constructed a phylogenetic tree of the DSCAM family based on full-length amino acid sequences (Figure 1). The tree revealed that DSCAM and DSCAML1 form two distinct clades that diverged after the split from sea urchin, suggesting that the gene duplication occurred early in the vertebrate lineage. Notably, all cyclostome orthologs were grouped within the DSCAML1 clade with strong bootstrap support (85%). This topology was also recovered in a phylogenetic tree constructed using the Bayesian approach (Supplementary Figure S1). Together, these results support the assignment of the cyclostome orthologs to *Dscaml1*.

**Figure 1.**
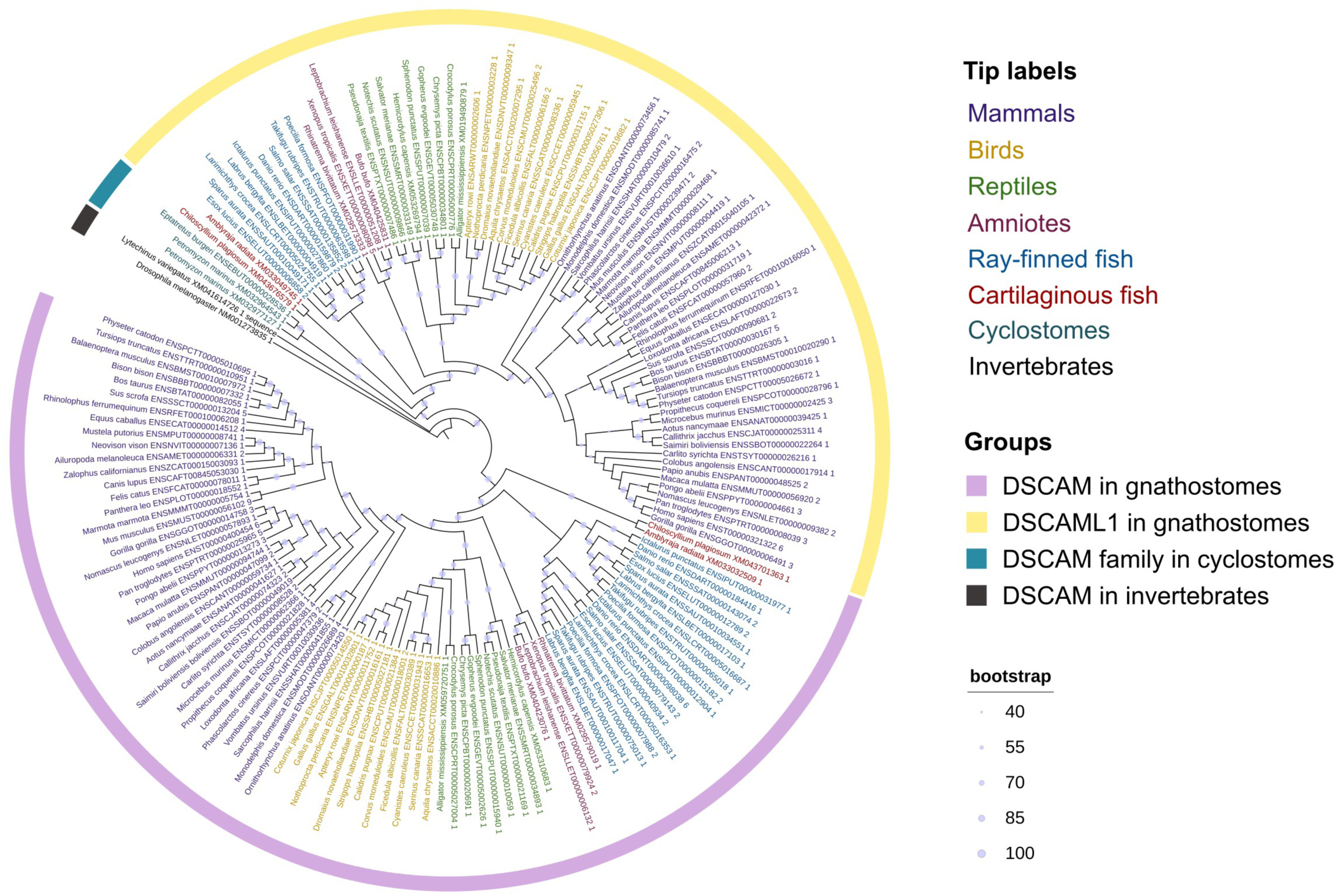
Phylogenetic relationship among DSCAM family in vertebrates. The phylogenetic tree of the vertebrate DSCAM family was constructed using the maximum likelihood method (IQ-Tree2). The tip labels are color coded according to their taxonomic classification. The ortholog DSCAM from *Drosophila melanogaster* was used as outgroup. The tree was edited and visualized by the iTOL v7. Ultrafast bootstrap values have been shown on each node.

We further examined the genomic organization of the *Dscam* and *Dscaml1* genes across vertebrates. While the local synteny around *Dscam* and *Dscaml1* is generally conserved among gnathostomes, the genomic context differs between gnathostomes and cyclostomes (Figure 2A,B). In gnathostomes, *Igsf5*, *Pcp4*, *Bace2*, and *Fam3b* are adjacent to the *Dscam* (Figure 2A); however, this synteny was not observed around *Dscam* family orthologs in cyclostomes (Figure 2B). In contrast, the *Dscaml1* locus shows a partially conserved synteny between gnathostomes and cyclostomes. For example, *Cep164* is located next to the *Dscaml1* in gnathostomes (Figure 2A) and is similarly positioned near one of the lamprey orthologs (Figure 2B). Additionally, *Htr3a*, *Zbtb16*, and *Ube4a*, which flank *Dscaml1* in gnathostomes (Figure 2A), are also found surrounding this ortholog in lamprey (Figure 2B). In hagfish, *Ube4a* is positioned near the *Dscam* family ortholog, and *Tlcd5*, which is also close to human *DSCAML1*, is located next to this ortholog (Figure 2B). These findings support the notion that these cyclostome orthologs correspond to *Dscaml1*. Lamprey possesses another *Dscam* family ortholog that lacks clear synteny with either *Dscam* or *Dscaml1* (Figure 2B).

**Figure 2.**
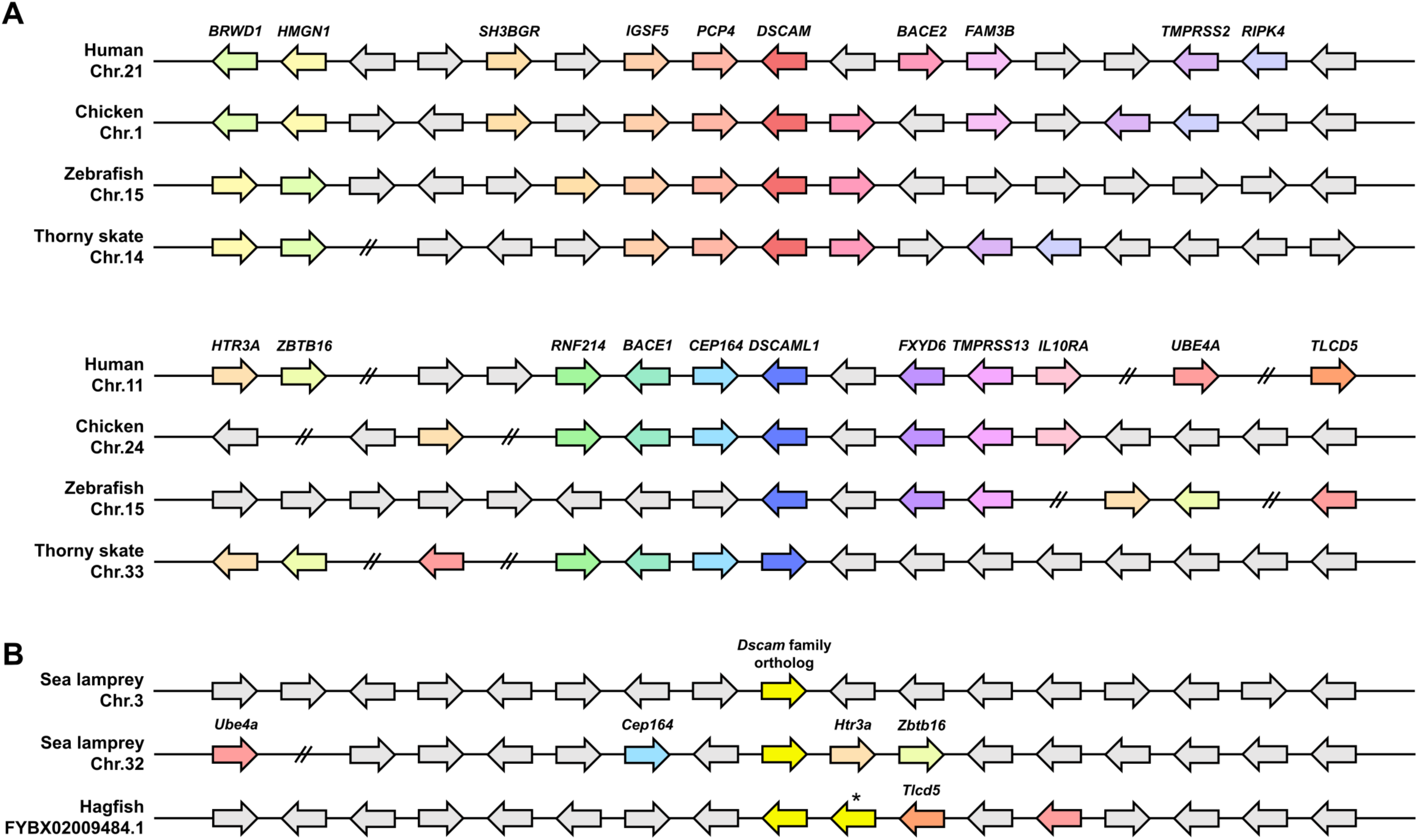
Synteny analysis of *Dscam* gene family in vertebrates. (**A**) Schematic representation of genes located upstream and downstream of *Dscam* family genes in osteichthyans. Top: the *Dscam* locus in osteichthyans. Bottom: the *Dscaml1* locus in osteichthyans. (**B**) Syntemy of *Dscam* family orthologs in cyclostomes. Arrows represent genes and their transcriptional orientations. Horizontal lines indicate chromosomes or sequence scaffolds, and distances are not drawn to scale. The asterisk indicates short fragments of *Dscam* family ortholog that may have arisen by partial duplication event in hagfish.

### Different evolutionary patterns between extracellular and intracellular domains of the *Dscam* gene family during gnathostome evolution

Given the apparent absence of *Dscam* in cyclostomes, we focused subsequent analyses on gnathostomes to characterize patterns of conservation and divergence in DSCAM family genes. A previous study suggested that sequence conservation between DSCAM and DSCAML1 is higher in the extracellular domain than in the intracellular domain (Agarwala et al., 2001). To evaluate this pattern, we calculated position-wise conservation scores using the AMAS method (Livingstone and Barton, 1993) from multiple alignments of gnathostome DSCAM and DSCAML1 orthologs (Supplementary Figure S2A). Consistent with this previous report, conservation scores were generally lower in the intracellular domains (Supplementary Figures S2A,B).

To assess differences in selective pressure acting on these domains, we calculated the ratio of nonsynonymous substitutions per nonsynonymous site (dN) to synonymous substitutions per synonymous site (dS). We first computed pairwise dN/dS ratios from multiple codon alignments for each domain. Within DSCAM, the intracellular domain showed a significantly lower pairwise dN/dS ratio than the extracellular domain (Figure 3A). In contrast, the intracellular domain of DSCAML1 showed a slightly higher pairwise dN/dS ratio than the extracellular domain (Figure 3A). Between paralogs, the extracellular domains showed no significant difference in pairwise dN/dS, whereas the intracellular domain of DSCAML1 exhibited significantly higher dN/dS than that of DSCAM (Figure 3A). Together, these results suggest domain- and paralog-specific differences in selective constraints during gnathostome evolution.

**Figure 3.**
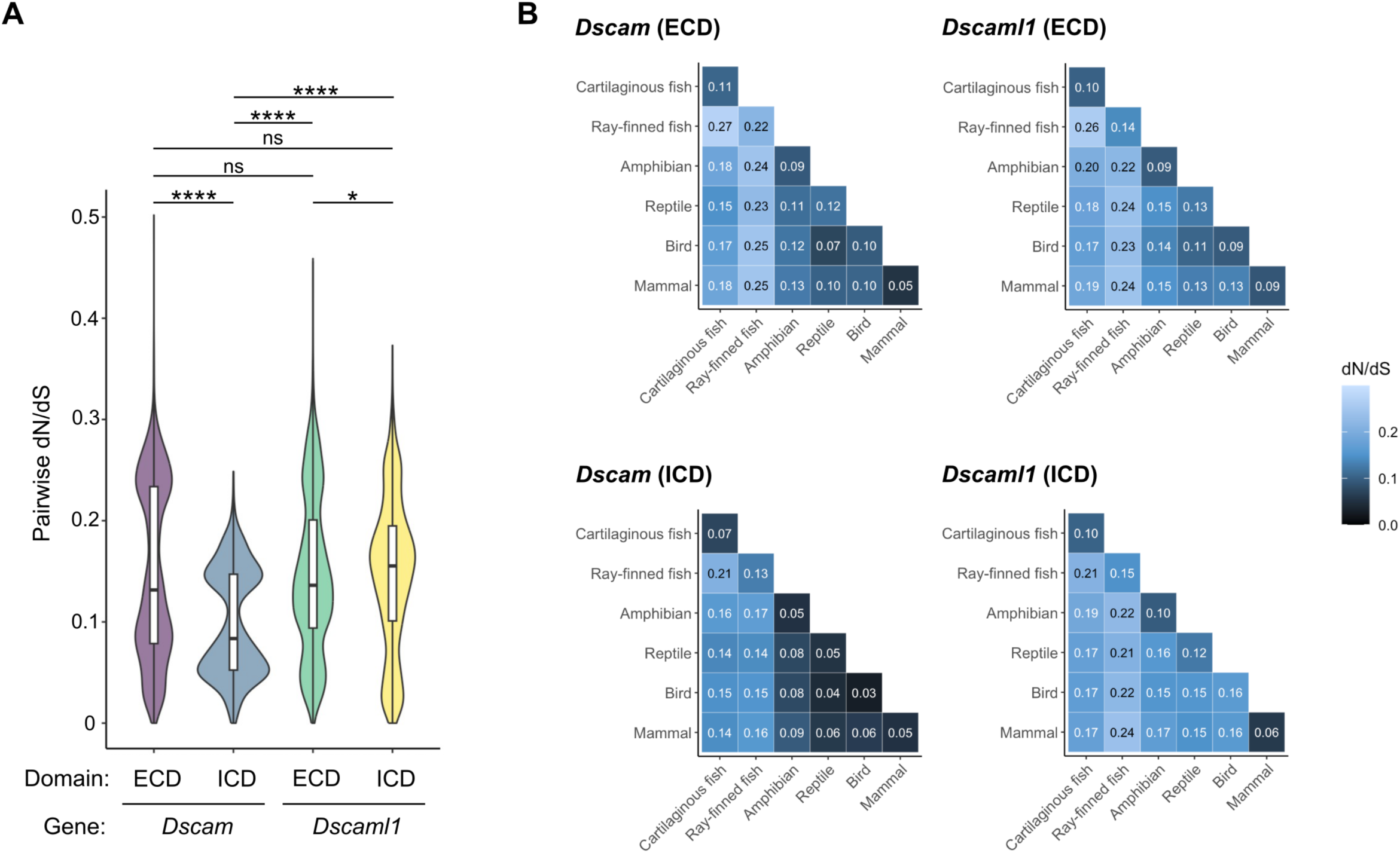
Pairwise dN/dS analysis of the extracellular and intracellular domains of the DSCAM family. **(A**) Pairwise dN/dS values were calculated for each domain of DSCAM and DSCAML1 orthologs from 74 gnathostomes. (**B**) The heatmap shows the average pairwise dN/dS ratios between clades. The values indicate the mean pairwise dN/dS ratios. *p*\* < 0.05 and *p*\*\*\*\* < 0.0001 (Dunn test).

We next compared pairwise dN/dS ratios across major vertebrate clades, including cartilaginous fishes, ray-finned fishes, amphibians, reptiles, birds, and mammals (Figure 3B). The extracellular domains of DSCAM and DSCAML1 showed similar patterns: average dN/dS ratios were lower in comparisons within tetrapods than in comparisons between more distantly related clades. For the intracellular domain of DSCAM, pairwise dN/dS ratios within tetrapods were markedly lower, indicating stronger purifying selection. In contrast, the intracellular domain of DSCAML1 exhibited generally higher pairwise dN/dS ratios than that of DSCAM within tetrapods, except among mammals. These results suggest that different evolutionary pressures have acted on the intracellular domains of DSCAM and DSCAML1 following the divergence of ray-finned fishes and tetrapods, while both domains in mammals have been subject to strong purifying selection.

We next tested whether the overall intensity of selection acting on the extracellular and intracellular domains of DSCAM and DSCAML1 changed after the split between ray-finned fishes and tetrapods using RELAX (Wertheim et al., 2015). We designed the tetrapod stem branch and descendant branches excluding mammals as the test set (non-mammal tetrapods) and used ray-finned fishes together with cartilaginous fishes as the reference set. For the extracellular domain, RELAX detected a significant intensification of selection (k > 1) on the test branches for both paralogs (Table 1), consistent with stronger evolutionary constraint in non-mammalian tetrapods.

**Table 1.** RELAX analysis for each domain of the DSCAM family.

| Gene | Domain | k<sup>a</sup> | 2ΔlnL | p-value<sup>b</sup> | Selection shift |
| --- | --- | --- | --- | --- | --- |
| *Dscam* | ECD | 2.29 | 177.32 | < 0.0001 | Intensification |
| *Dscam* | ICD | 1.17 | 9.68 | < 0.01 | Intensification |
| *Dscaml1* | ECD | 1.97 | 86.23 | < 0.0001 | Intensification |
| *Dscaml1* | ICD | 1.09 | 2.10 | 0.147 | - |

<sup>a</sup> Selection intensity parameter. k > 1 indicates that selection strength has been intensified along the test branches.
<sup>b</sup> Likelihood ratio test p-values.

For the intracellular domain, RELAX similarly detected a significant intensification of selection for DSCAM on the test branches (Table 1), whereas no significant change in selection intensity was detected for DSCAML1 (Table 1). Together with the pairwise dN/dS patterns, these results suggest that the intracellular domain of DSCAML1 did not undergo the same reinforcement of constraint observed for DSCAM during tetrapod evolution.

Given that RELAX analyses indicated differential reinforcement of selection between domains and paralogs, we next asked whether a subset of sites might have experienced episodic shifts in selective pressure in specific lineages using PAML’s branch-site model A test. We focused on the stem branches of osteichthyes, tetrapods, amniotes, and mammals as foreground branches. Likelihood-ratio tests (LRTs) did not provide significant evidence for episodic positive selection in DSCAM in any lineage or domain (Table 2). In contrast, LRTs supported episodic positive selection in DSCAML1, with significant signals detected in the extracellular domain across multiple forground lineages and in the intracellular domain on the tetrapod stem branch (Table 2).

**Table 2.** Branch-site model applied to the stem branch of each major clade.

| Gene | Domain | Foreground branch | Proportion of site class | $\omega$ | 2ΔlnL | LRT $p$-value<sup>a</sup> | Possible positive selection sites (BEB posterior probability)<sup>b</sup> |
| --- | --- | --- | --- | --- | --- | --- | --- |
| *Dscam* | ECD | Osteichthyes | 0: 0.94788<br>1: 0.00409<br>2a: 0.04782<br>2b: 0.00021 | $\omega_0$ = 0.04082<br>$\omega_1$ = 1.00000<br>$\omega_2$ = 2.17157 | 2.77346 | 0.0958 | 65K (0.919), 114R (0.977), 176S (0.991), 655R (0.964), 682L (0.957), 818Q (0.992), 821H (0.984), 977P (0.970), 1101S (0.963) |
| | | Tetrapod | 0: 0.99020<br>1: 0.00412<br>2a: 0.00566<br>2b: 0.00002 | $\omega_0$ = 0.04122<br>$\omega_1$ = 1.00000<br>$\omega_2$ = 7.61108 | 3.305312 | 0.0691 | 251A (0.978), 288M (0.959) |
| | | Amniote | 0: 0.99581<br>1: 0.00418<br>2a: 0.00000<br>2b: 0.00000 | $\omega_0$ = 0.04135<br>$\omega_1$ = 1.00000<br>$\omega_2$ = 1.00000 | 0 | 1 | |
| | | Mammal | 0: 0.94247<br>1: 0.00403<br>2a: 0.05328<br>2b: 0.00023 | $\omega_0$ = 0.04078<br>$\omega_1$ = 1.00000<br>$\omega_2$ = 2.73582 | 2.45653 | 0.117 | 128R (0.987), 204H (0.990), 773S (0.992) |
| | ICD | Osteichthyes | 0: 0.87655<br>1: 0.01734<br>2a: 0.10405<br>2b: 0.00206 | $\omega_0$ = 0.02681<br>$\omega_1$ = 1.00000<br>$\omega_2$ = 999.00000 | 0.5321 | 0.468 | |
| | | Tetrapod | 0: 0.96465<br>1: 0.02058<br>2a: 0.01446<br>2b: 0.00031 | $\omega_0$ = 0.02700<br>$\omega_1$ = 1.00000<br>$\omega_2$ = 8.40784 | 3.1016 | 0.0782 | 340Q (0.983) |
| | | Amniote | 0: 0.00000<br>1: 0.00000<br>2a: 0.97936<br>2b: 0.02064 | $\omega_0$ = 0.02709<br>$\omega_1$ = 1.00000<br>$\omega_2$ = 1.00000 | 0 | 1 | |
| | | Mammal | 0: 0.97884<br>1: 0.02116<br>2a: 0.00000<br>2b: 0.00000 | $\omega_0$ = 0.02736<br>$\omega_1$ = 1.00000<br>$\omega_2$ = 1.00000 | 0 | 1 | |
| *Dscaml1* | ECD | Osteichthyes | 0: 0.89089<br>1: 0.04774<br>2a: 0.05825<br>2b: 0.00312 | $\omega_0$ = 0.04009<br>$\omega_1$ = 1.00000<br>$\omega_2$ = 3.65686 | 15.5956 | < 0.001 | 80V (0.979), 102V (0.955), 123G (0.959), 137A (0.974), 234F (0.961), 298A (0.932), 322G (0.959), 396A (0.975), 738S (0.961), 893V (0.960), 977T (0.973), 995S (0.906), 1000S (0.989), 1136S (0.987), 1167V (0.959), 1169S (0.960), 1172E (0.989), 1208R (0.973), 1209E (0.904), 1239S (0.998), 1303S (0.986), 1306Q (0.983), 1317G (0.945), 1337E (0.925), 1367R (0.985), 1394S (0.993), 1598T (0.985), 1641A (0.961) |
| | | Tetrapod | 0: 0.90705<br>1: 0.04822<br>2a: 0.04247<br>2b: 0.00226 | $\omega_0$ = 0.04093<br>$\omega_1$ = 1.00000<br>$\omega_2$ = 999.00000 | 5.4765 | < 0.05 | |
| | | Amniote | 0: 0.94650<br>1: 0.05042<br>2a: 0.00292<br>2b: 0.00016 | $\omega_0$ = 0.04127<br>$\omega_1$ = 1.00000<br>$\omega_2$ = 27.90127 | 6.2151 | < 0.05 | |
| | | Mammal | 0: 0.94700<br>1: 0.05033<br>2a: 0.00253<br>2b: 0.00013 | $\omega_0$ = 0.04128<br>$\omega_1$ = 1.00000<br>$\omega_2$ = 11.40634 | 5.77672 | < 0.05 | 70D (0.942), 893V (0.941), 925S (0.952) |
| | ICD | Osteichthyes | 0: 0.86154<br>1: 0.09643<br>2a: 0.03780<br>2b: 0.00423 | $\omega_0$ = 0.03077<br>$\omega_1$ = 1.00000<br>$\omega_2$ = 1.49311 | 0.31678 | 0.574 | 267A (0.929), 337A (0.901) |
| | | Tetrapod | 0: 0.86532<br>1: 0.09719<br>2a: 0.03371<br>2b: 0.00379 | $\omega_0$ = 0.03077<br>$\omega_1$ = 1.00000<br>$\omega_2$ = 999.00000 | 8.04292 | < 0.01 | 80A (0.918), 296E (0.921), 329A (0.965), 332L (0.958), 342G (0.939), 346P (0.91), 423V (0.942) |
| | | Amniote | 0: 0.78405<br>1: 0.09038<br>2a: 0.11260<br>2b: 0.01298 | $\omega_0$ = 0.03106<br>$\omega_1$ = 1.00000<br>$\omega_2$ = 2.04509 | 0 | 1 | |
| | | Mammal | 0: 0.89664<br>1: 0.10336<br>2a: 0.00000<br>2b: 0.00000 | $\omega_0$ = 0.03106<br>$\omega_1$ = 1.00000<br>$\omega_2$ = 1.00000 | 0 | 1 | |

<sup>a</sup> Likelihood ratio test $p$ -values. <sup>b</sup> Sites potentially under positive selection based on BEB analysis. Sites with a posterior probability > 0.90 are considered candidates for selection. Site positions are numbered from the N-terminus for ECD and from immediately after the transmembrane domain for ICD.

Despite the lack of significant LRT support, Bayes Empirical Bayes (BEB) analysis highlighted 14 candidate sites in the extracellular domain and one site in the intracellular domain of DSCAM with posterior probabilities > 0.90 (Table 2). For DSCAML1, BEB analysis identified 30 positively selected sites in the extracellular domain, 28 of which were assigned to the osteichthyan ancestor, and nine sites in the intracellular domain, including seven sites in the tetrapod ancestor (Table 2). Taken together, these results indicate distinct lineage- and site-specific patterns of selective shifts between DSCAM and DSCAML1.

### Functional divergence of the intracellular domain between DSCAM and DSCAML1

Given the distinct evolutionary patterns observed for the intracellular domains of DSCAM and DSCAML1, we next asked whether these intracellular domains exhibit signatures consistent with functional divergence during tetrapod evolution. Using DIVERGE v3.0 (Gu et al., 2013) and amino acid sequences from 12 tetrapod species, we detected significant Type I functional divergence between DSCAM and DSCAML1 for the intracellular domain (θ > 0; Table 3), supporting a shift in site-specific evolutionary constraints between the paralogs. Type II analysis likewise supported divergence for the ICD, consistent with shifts in amino acid physicochemical properties between the paralogs (Table 3). A similar pattern was also observed for the extracellular domain (Table 3).

**Table 3.** Functional divergence of each domain of the DSCAM family.

| Clade 1 | Clade 2 | Type I |  |  | Type II |  |  |
| --- | --- | --- | --- | --- | --- | --- | --- |
|  |  | MFE Theta | MFE se | I: *p*-value<sup>a</sup> | Theta-II | Theta SE | II: *p*-value<sup>a</sup> |
| DSCAM-ECD | DSCAML1-ECD | 0.557342 | 0.056026 | < 0.0001 | 0.266351 | 0.021056 | < 0.0001 |
| DSCAM-ICD | DSCAML1-ICD | 0.142066 | 0.039115 | < 0.001 | 0.358524 | 0.037526 | < 0.0001 |

<sup>a</sup> *P*-values evaluated by calculating the Z-score ( $\theta/SE$ ) under the assumption of a normal distribution.

To further explore potential functional differences in the intracellular domains, we referred to the AlphaFold Protein Structure Database (Varadi et al., 2024) and used ColabFold (Mirdita et al., 2022) to predict protein structures for DSCAM and DSCAML1 (Figures 4A,B). However, the AlphaFold2-predicted models showed very low confidence scores (pLDDT) across the intracellular regions in both proteins in the available models and in our ColabFold predictions. This suggests that the intracellular domains are likely composed primarily of intrinsically disordered regions (IDRs) (Akdel et al., 2022). Indeed, IUPred3 (Erdős et al., 2021) predicted that nearly the entire intracellular domain of both proteins is disordered (Figure 4C).

**Figure 4.**
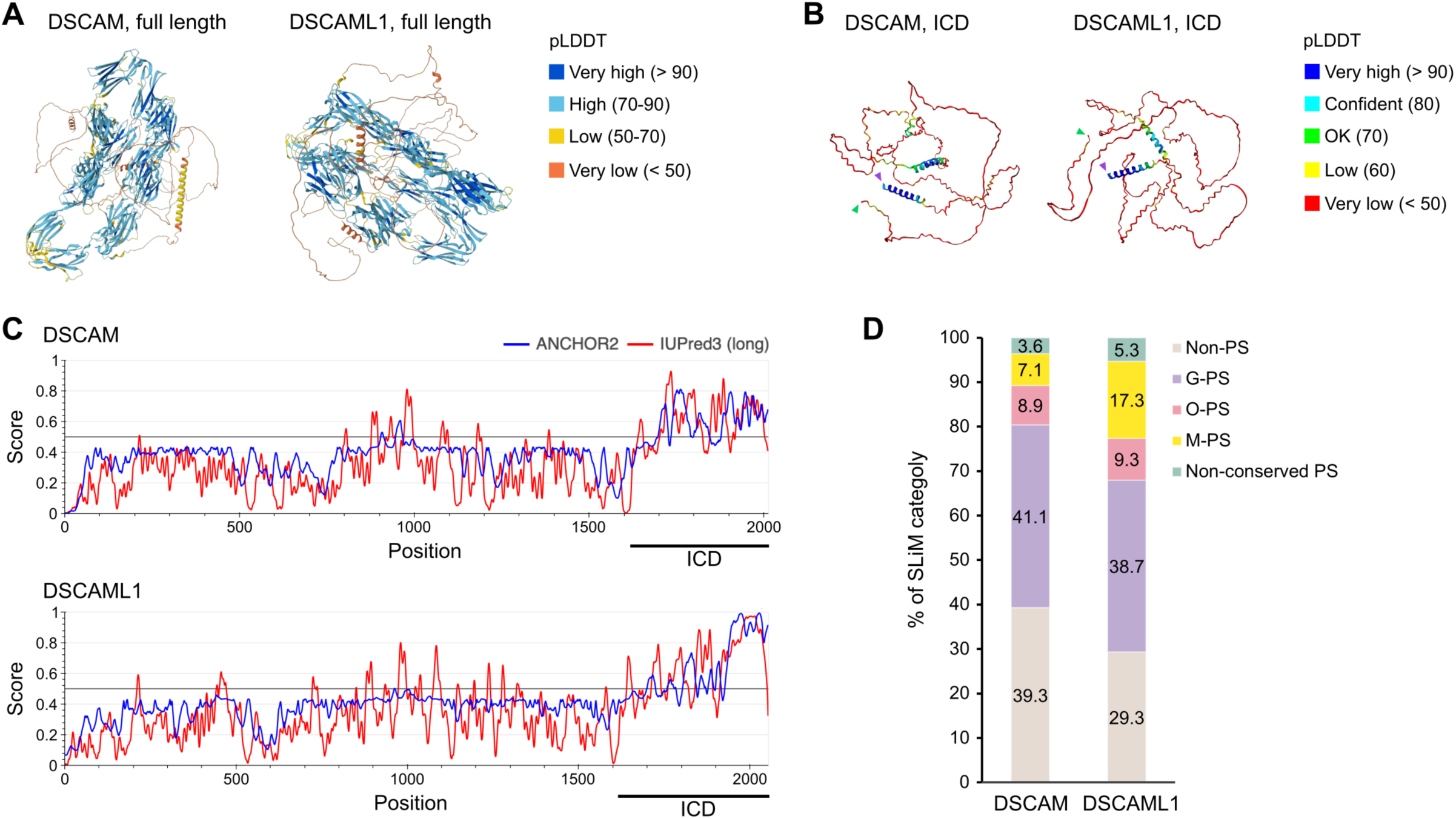
Properties of the intracellular domains of the DSCAM family. (**A**) Predicted structures of human DSCAM (left) and DSCAML1 (right) generated from Alphafold2 (AlphaFold DB: AF-O60469-F1-v4 and AF-Q8TD84-F1-v4, respectively). The structures are colored by pLDDT (confidence in local structure) score. (**B**) Predicted structures of intracellular domains of human DSCAM (left) and DSCAML1 (right) generated from Alphafold2 (using ColabFold). Purple arrowheads and green arrowheads represent N-terminus and C-terminus of the intracellular domain, respectively. (**C**) Prediction of the degree of the disorder in DSCAM (top) and DSCAML1 (bottom). IUPred3 (red line), which predicts intrinsically disordered regions in a given amino acid sequence, and ANCHOR2 (blue line), which predicts the disordered segments that can interact with globular proteins. A region with an IUPred3 score continuously greater than 0.5 indicates a disordered region. A region with ANCHOR2 score continuously greater than 0.5 in a disordered region indicates a disordered segment that can interact with globular proteins. The region of the intracellular domain is shown below the plot, respectively. (**D**) Ratio of SLiMs in the intracellular domains of DSCAM and DSCAML1 in human. SLiMs were categorized into two groups: conserved SLiMs and paralog-specific SLiMs. Paralog-specific SLiMs were further divided into four subgroups based on interspecies comparison. (1) gnathostome-conserved paralog-specific SLiMs (G-PS), (2) osteichthyan-conserved paralog-specific SLiMs (O-PS), (3) mammal-conserved paralog-specific SLiMs (M-PS), and (4) non-conserved paralog-specific SLiMs (non-conserved PS). The proportion of SLiMs in each category is shown in the graph.

IDRs play crucial roles in the regulation of signaling pathways and key cellular processes (Wright and Dyson, 2015). These functions are frequently mediated by short linear motifs (SLiMs), which are 3−10 amino acid sequences enriched within IDRs (Davey et al., 2012). Due to their low-affinity and reversible binding properties, SLiMs are particularly well-suited for transient protein–protein interactions and dynamic regulation. To further investigate possible functional divergence, we compared SLiMs associated with protein–protein interactions (i.e., classes LIG and DOC) within the intracellular domains of human DSCAM and DSCAML1. Using the ELM resource (Kumar et al., 2024), we predicted 56 SLiM instances in the intracellular domain of DSCAM and 75 in that of DSCAML1 (Supplementary Table S2). Among these, 34 SLiMs (60.7%) in DSCAM and 53 in DSCAML1 (70.7%) were not conserved at corresponding positions in the alignment between the paralogs and were considered paralog-specific (Figure 4D; Supplementary Table S3).

To examine the evolutionary conservation of paralog-specific SLiMs, we compared them with SLiMs identified in the intracellular domains of DSCAM and DSCAML1 orthologs in mouse, zebrafish and thorny skate (Supplementary Tables S4-S6). We subcategorized the human paralog-specific SLiMs into four groups: (1) gnathostome-conserved paralog-specific SLiMs (G-PS), (2) osteichthyan-conserved paralog-specific SLiMs (O-PS), (3) mammal-conserved paralog-specific SLiMs (M-PS), and (4) non-conserved paralog-specific SLiMs (non-conserved PS) (see Methods; Supplementary Table S7). The proportions of G-PS were comparable between the two genes: 23 SLiMs (41.1%) in DSCAM and 29 (38.7%) in DSCAML1 (Figure 4D). A similar pattern was observed for O-PS: 5 (8.9%) in DSCAM and 7 (9.3%) in DSCAML1 and non-conserved PS: 2 (3.6%) in DSCAM and 4 (5.3%) in DSCAML1 (Figure 4D). In contrast, M-PS showed an asymmetric distribution, with 4 (7.1%) in DSCAM and 13 (17.3%) in DSCAML1 (Figure 4D). Notably, six of the nine amino acid sites predicted by PAML to have undergone positive selection in the intracellular domain of DSCAML1 were located within human paralog-specific SLiMs, including four sites within M-PS (Supplementary Table S8).

To further investigate functional consequences of divergence, we reanalyzed previously published bulk RNA-seq data from HEK293 cells stably expressing the intracellular domains of either gene (Sachse et al., 2019). DEGs were identified by comparing expression profiles of DSCAM- or DSCAML1-expressing cells to control cells. DSCAM expression resulted in 911 upregulated and 407 downregulated genes, while DSCAML1 expression led to 624 upregulated and 521 downregulated genes (Figure 5A). Gene Ontology (GO) enrichment analysis revealed that DEGs from DSCAM-expressing cells were associated with 274 biological process terms in ShinyGO and 127 terms in PANTHER (FDR < 0.05; Figure 5B). In contrast, DSCAML1-expressing cells showed enrichment for 955 GO terms in ShinyGO and 480 in PANTHER. In both datasets, we observed enrichment in developmental processes such as nervous system development and neuron projection development (Supplementary Figures S3A,B), consistent with previous findings (Sachse et al., 2019).

**Figure 5.**
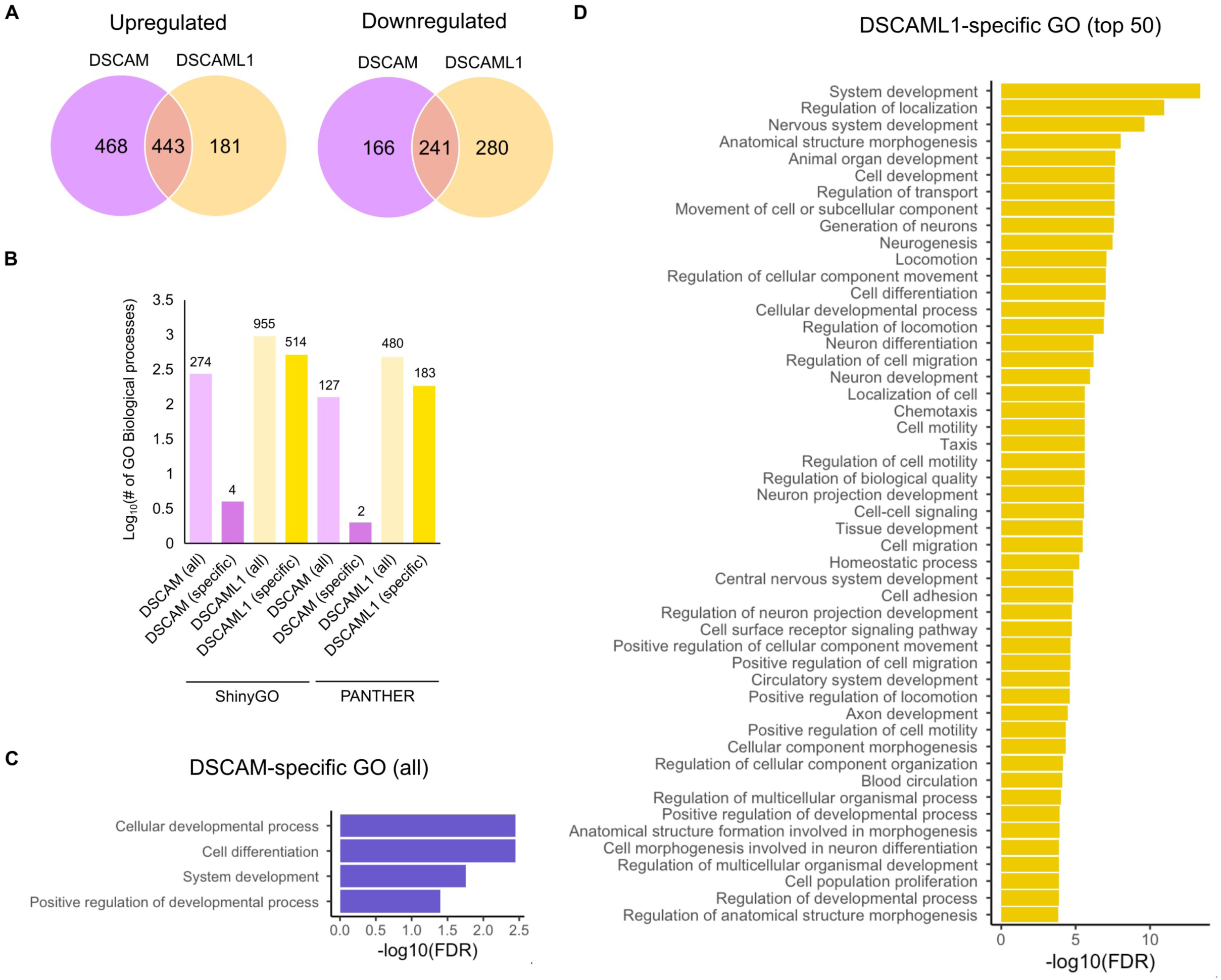
Transcriptomic differences by overexpression of the intracellular domain of DSCAM and DSCAML1. (**A**) Venn diagrams showing the number of overlapping and individual DEGs between DSCAM ICD and DSCAML1 ICD. Left: upregulated genes by the overexpression, right: downregulated genes by the overexpression. (**B**) Number of GO terms related to biological process. GO terms by whole DEGs (all) and specific DEGs (specific) are shown. GO analysis were performed using two tools; ShinyGO and PANTHER. The numbers of GO terms above the bar graph. (**C**) The result of GO analysis performed with ShinyGO. GO terms (biological process) in DSCAM-specific DEGs (top) and DSCAML1-specific DEGs (bottom) are shown. All and top 50 GO terms that are significantly enriched (FDR < 0.05) are shown for DSCAM and DSCAML1, respectively.

To identify paralog-specific gene regulation, we compared DEG sets. DSCAM-specific regulation included 468 upregulated and 166 downregulated genes, whereas DSCAML1-specific regulation comprised 181 upregulated and 280 downregulated genes (Figure 5A). GO analysis of DSCAM-specific DEGs identified four biological process terms in ShinyGO and two in PANTHER, all related to developmental processes (Figures 5B,C; Supplementary Figure S4A). In contrast, DSCAML1-specific DEGs were associated with 514 terms in ShinyGO and 183 in PANTHER, enriched for functions such as neurogenesis, neuron projection development, cell migration, signaling pathways, and cell proliferation (Figure 5C; Supplementary Figure S4B). The larger number and broader range of GO terms associated with DSCAML1-specific DEGs suggest that DSCAML1 affects a wider spectrum of biological processes under these experimental conditions.

## Discussion

In arthropods, DSCAM has evolved to function in neuronal cell-surface recognition, thereby contributing to the establishment of proper neural networks. In contrast, this function has independently emerged in vertebrates through the convergent evolution of cPcdhs (Dong et al., 2023). Despite lacking the extensive isoform diversity seen in arthropods, the DSCAM family in vertebrates is also highly conserved and has been implicated in neural circuit formation and brain function. Thus, the DSCAM family and cPcdhs represent a compelling model of convergent evolution in neural systems. To date, however, the evolutionary history of the vertebrate DSCAM family remains poorly understood, as most phylogenetic studies have focused on invertebrate DSCAM to explore the diversification of the extracellular domain (Crayton et al., 2006; Brites et al., 2008; Lee et al., 2010; Cao et al., 2018; Wiseglass and Rubinstein, 2024). Moreover, no previous study has investigated the functional divergence between DSCAM and DSCAML1 from a molecular evolutionary perspective. In the present study, we elucidated the phylogenetic relationships of DSCAM and DSCAML1 across vertebrates and examined the evolutionary pressures acting separately on their extracellular and intracellular domains. Our findings reveal distinct evolutionary trajectories for DSCAM and DSCAML1, and suggest functional divergence between their intracellular domains. These results imply that the evolution of vertebrate neural networks is shaped by a complex interplay of functional divergence, molecular innovation, and historical contingency.

Consistent with previous research indicating that the duplication of the *Dscam* gene occurred prior to vertebrate diversification (Crayton et al., 2006), our study supports the view that this duplication event likely took place after the divergence of echinoderms and chordates. However, its precise timing remains unresolved for two main reasons. First, *Dscam* orthologs were not detected in cephalochordates or urochordates, consistent with the earlier report (Brites et al., 2008). Second, the orthologous relationship between cyclostome and gnathostome genes could not be confidently determined. Cyclostome genomes exhibit nucleotide composition biases and have undergone a lineage-specific whole-genome duplication (Nakatani et al., 2021; Yu et al., 2024), both of which complicate orthology inference. Nevertheless, synteny analysis revealed the presence of *Dscaml1* in cyclostomes, and both ML and Bayesian phylogenetic trees placed cyclostome *Dscaml1* together with gnathostome *Dscaml1*, following the divergence between *Dscam* and *Dscaml1*. These observations suggest that the gene duplication event likely occurred before the split of cyclostomes and gnathostomes. In lamprey, two *Dscam* family orthologs were identified; one of them lacks the conserved synteny shared with gnathostome *Dscam* and *Dscaml1*, yet clusters within the *Dscaml1* clade in phylogenetic analyses. Two possible evolutionary scenarios could explain these results. (1) The common ancestor of cyclostomes may have lost *Dscam*, while a subsequent whole-genome duplication within the cyclostome lineage generated a duplicate of *Dscaml1*, with the hagfish lineage later losing one of the copies. (2) Alternatively, hagfish may have lost *Dscam* after diverging from the lamprey lineage, and the rapid evolution of lamprey *Dscam* may have obscured its placement in the phylogenetic tree, leading to misassignment.

Protein evolutionary rates are associated with protein localization and exposure to the extracellular environment (Julenius and Pedersen, 2006; Liao et al., 2010). In transmembrane proteins, extracellular and intracellular domains often experience distinct selective constraints, with extracellular regions frequently evolving faster than cytoplasmic regions. Consistent with this pattern, the intracellular domain of DSCAM exhibited lower dN/dS ratios than its extracellular domain, with especially low values among tetrapods. By contrast, the intracellular domain of DSCAML1 showed relatively higher dN/dS ratios, suggesting that the two paralogs have experienced distinct selective regimes on their intracellular domains. Specifically, our results indicate that selection intensity on the DSCAM intracellular domain increased after the emergence of tetrapods, whereas no comparable shift was detected for the DSCAML1 intracellular domain. A weaker reinforcement of constraint on the DSCAML1 intracellular domain may have permitted the accumulation of lineage-specific amino acid substitutions, potentially contributing to functional divergence between DSCAM and DSCAML1. Consistent with this interpretation, branch-site analyses in PAML further suggested episodic, site-specific shifts in selective pressure acting on the DSCAML1 intracellular domain. In mammals, the consistently low dN/dS ratios for the DSCAML1 intracellular domain suggest renewed or stronger functional constraint, implying that this domain later became more strongly constrained, potentially reflecting stabilization of an essential function after earlier divergence. This is consistent with previous studies showing that genes involved in nervous system functions tend to be subject to stronger purifying selection in mammals (Brawand et al., 2011; Kryuchkova-Mostacci and Robinson-Rechavi, 2015).

Although the extracellular domains of DSCAM and DSCAML1 are similarly constrained overall, the branch-site analyses indicate that the extracellular domain of DSCAML1 experienced more pronounced episodic, site-specific adaptive changes than that of DSCAM, particularly along the osteichthyan stem lineage. Such localized signals are difficult to detect using pairwise dN/dS, which averages substitution rates across the entire domain, but can be captured by branch-site models that are sensitive to lineage- and site-specific effects. Most inferred sites fall within immunoglobulin-like and fibronectin type III domains, suggesting that these substitutions may have affected interaction-partner specificity or adhesion properties. Indeed, DSCAM and DSCAML1 exhibit different adhesion behaviors at cell–cell interfaces (Guo et al., 2021). One possibility is that early changes in extracellular adhesion altered the functional context of the protein, thereby creating evolutionary opportunity for subsequent shifts in selective constraint on the intracellular domain. Further experimental work will be required to determine how these candidate positively selected substitutions affect the molecular functions and biological roles of DSCAML1.

Previous studies have demonstrated that both mammalian DSCAM and DSCAML1 are involved in key neurodevelopmental processes, including self-avoidance, neurite development, and neuronal migration (Fuerst et al., 2009; Zhang et al., 2015; Sachse et al., 2019; Mitsogiannis et al., 2020). As shown in Figure 3B, the intracellular domain of mammalian DSCAML1 shows even lower dN/dS ratios compared to other vertebrate lineages, suggesting that this region has been subject to stronger functional constraints during mammalian evolution. This increased constraint likely reflects its critical biological role following functional divergence. However, the extent to which these paralogs have diverged in function following gene duplication remains poorly understood. In the present study, we indicate potential functional divergence between mammalian DSCAM and DSCAML1, with a particular focus on their intracellular domains. The majority of intracellular domains of DSCAM and DSCAML1 are predicted to consist of IDRs, where gain or loss of SLiMs throughout the evolution can lead to alteration of protein function and contribute to functional divergence after gene duplication (Davey et al., 2015). Due to their low-affinity and reversible binding properties, SLiMs are particularly well-suited for transient protein–protein interactions and dynamic regulation. Consequently, they contribute significantly to the assembly and modulation of protein interaction networks and are central to coordinating complex intracellular signaling pathways (Davey et al., 2012; Van Roey et al., 2014). Our comparative analysis of SLiMs in the intracellular domains of mammalian DSCAM and DSCAML1 indicates that the two paralogs harbor distinct motif repertoires, which likely confer differences in their potential protein interaction partners. For instance, only DSCAML1 possesses class I and class II SH3 domain-binding motifs, and the number of non-canonical class I SH3-binding motifs is expanded in DSCAML1 relative to DSCAM. These motifs are known to mediate interactions with SH3 domain-containing adaptor proteins, which play crucial roles in diverse cellular processes, including cytoskeletal dynamics, migration, endocytosis, apoptosis regulation, and proteasomal degradation (Mehrabipour et al., 2023). The enrichment of these motifs in DSCAML1 implies that it may have evolved an expanded role in SH3-mediated signaling networks. Importantly, DSCAML1 harbors a greater number of M-PS compared to DSCAM, indicating an asymmetric pattern of motif acquisition. This difference may reflect distinct evolutionary trajectories: DSCAM appears to have retained ancestral functions under strong purifying selection, whereas DSCAML1 may have undergone shifts in selective pressure during tetrapod evolution. Supporting this notion, several amino acid sites predicted to be under positive selection in DSCAML1 are located within paralog-specific SLiMs. In some cases, these substitutions occur at functionally conserved residues within SLiMs—for example, the leucine in the RxL motif (a Cyclin/CDK recognition site), the proline in the xxx[PV]xxP motif (recognized by SH3 domains), and the valine in the xx(T)xx[ILV]x motif (recognized by FHA domains). Such modifications may have altered the binding specificity or regulatory potential of these motifs, contributing to the functional innovation of DSCAML1 during vertebrate evolution.

Further evidence for functional divergence comes from transcriptomic data. Expression of the intracellular domains of the DSCAM family in HEK293 cells revealed largely non-overlapping sets of DEGs. While the numbers of paralog-specific DEGs were comparable, DSCAML1-specific DEGs were associated with a broader range of biological processes. Notably, only DSCAML1-specific DEGs, in addition to DSCAM/DSCAML1-common DEGs, were enriched for neural development-related pathways, suggesting that DSCAML1 may have acquired additional functions related to the nervous system development. Moreover, DSCAML1-specific DEGs were enriched for genes involved in cell migration, which may be attributed to the abundance of SH3 domain–binding motifs in the intracellular domain of DSCAML1. Since SH3-containing proteins are involved in cytoskeletal organization and its regulation (Mehrabipour et al., 2023), these motifs could mediate interactions that influence cellular motility. Together, these findings suggest that DSCAML1 has acquired the capacity to affect a wider array of cellular programs, consistent with the gain of novel functional elements such as paralog-specific SLiMs. It should be noted that this bulk RNA-seq analysis was conducted using a non-neuronal cell line. Therefore, future investigations using neuronal cell cultures or brain tissues will be necessary to fully elucidate the functional divergence between DSCAM and DSCAML1 in a more physiologically relevant context.

Overall, this study provides an evolutionary analysis of the DSCAM family in vertebrates, highlighting functional divergence in the intracellular domains. Our phylogenetic analyses suggest that the duplication event giving rise to DSCAM and DSCAML1 occurred prior to the divergence of cyclostomes and gnathostomes, likely after the split between echinoderms and chordates. DSCAM and DSCAML1 appear to have undergone different selective pressure in their intracellular domain throughout gnathostome evolution. Some amino acid changes in the intracellular domain may have contributed to functional innovation in certain vertebrate lineages; in mammals, our focal clade in the latter analyses, the consistently low pairwise dN/dS ratios indicate strong purifying selection, consistent with an essential role of this domain after functional divergence. Supporting this, expression of the intracellular domain of human DSCAML1 resulted in distinct gene expression changes compared to DSCAM. These findings suggest that the DSCAM family has evolved to play critical roles in shaping the complex neural networks of vertebrates, not through the expansion of combinatorial molecular codes as seen in arthropods, but through functional diversification following gene duplication.

## Author contributions

KH: Conceptualization, Analysis, Funding acquisition, Writing – original draft, Writing – review & editing. YW: Writing – review & editing. HO: Writing – review & editing. MH: Supervision, Funding acquisition, Writing – review & editing. All authors contributed to the article and approved the submitted version.

---

**Supplementary Figure S1.**
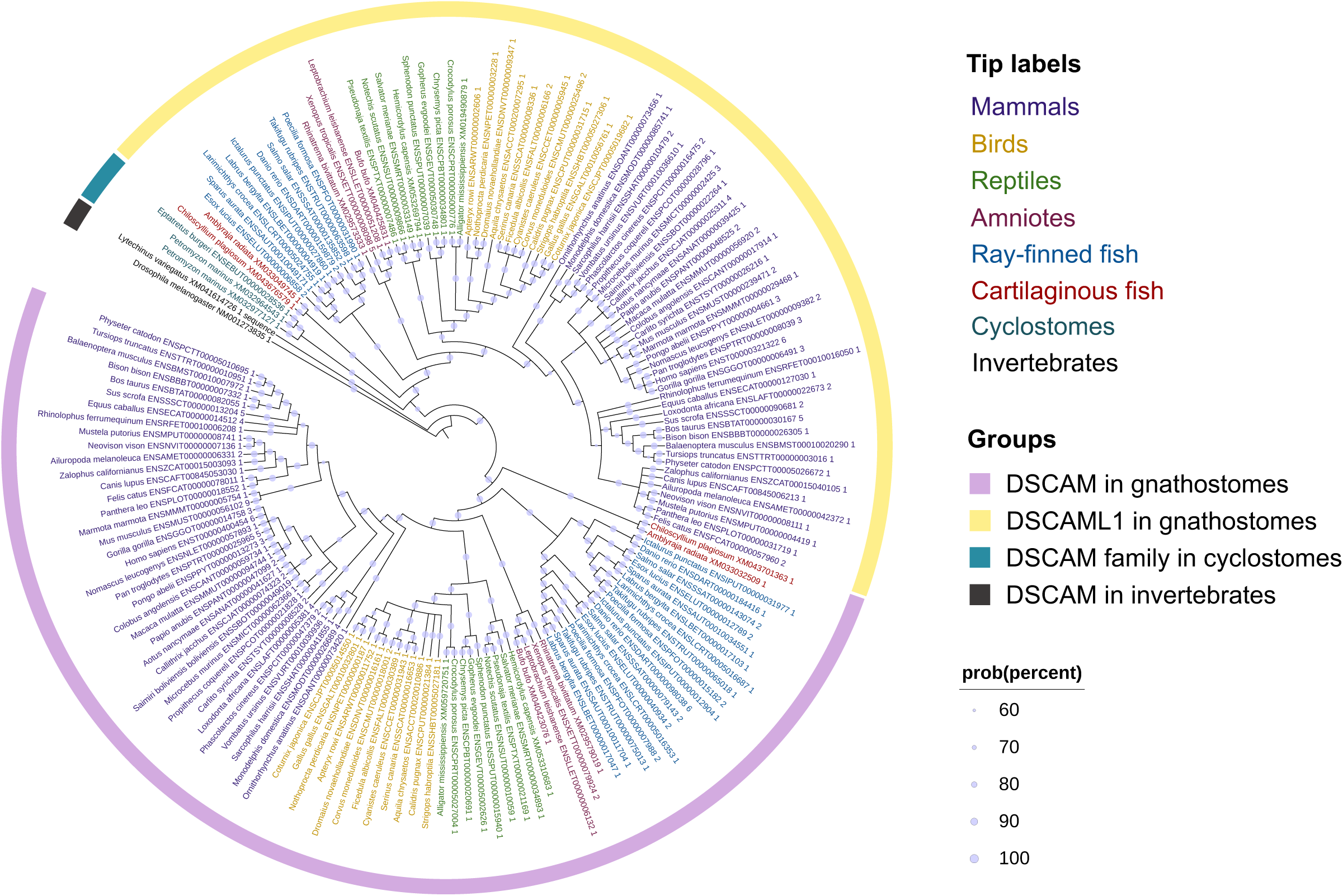
Bayesian phylogenetic tree of vertebrate DSCAM family. The phylogenetic tree of the vertebrate DSCAM family was constructed using the Bayesian method (MrBayes). The tip labels are color coded according to their taxonomic classification. The ortholog DSCAM from *Drosophila melanogaster* was used as outgroup. The tree was edited and visualized by the iTOL v7. Posterior probability values have been shown on each node.

**Supplementary Figure S2.**
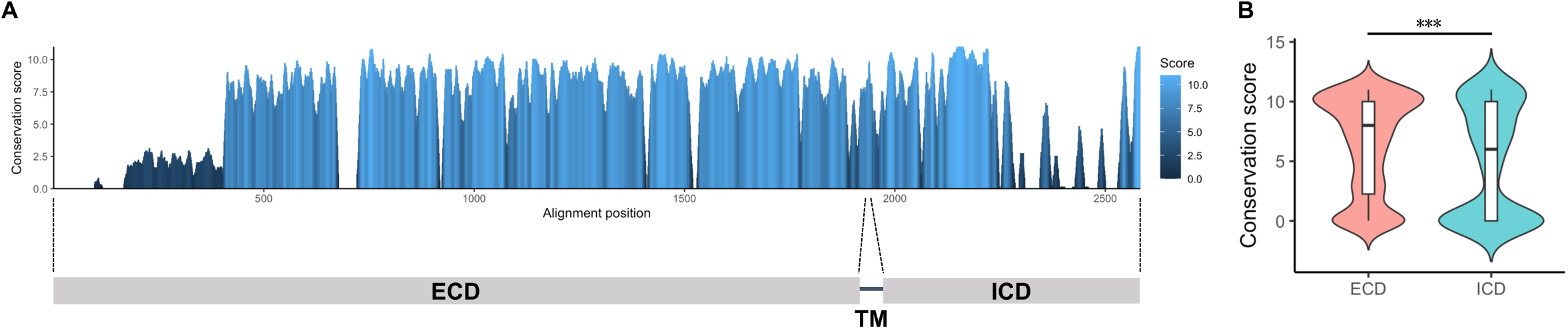
Amino acid conservation of gnathostome DSCAM family. (A) Top: Amino acid conservation score along an alignment of DSCAM family homologs calculated by the AMAS method (JalView). The histogram shows average of 10 amino acids window of conservation score for each position of the alignment. Bottom: Schematic diagram of DSCAM family protein. (B) The violin plot and box plot show the conservation score for each position of the alignment in the extracellular domain and intracellular domain. ECD, extracellular domain; TM, transmembrane domain; and ICD, intracellular domain. *p*\*\*\* < 0.001 (Wilcoxon rank sum test).

**Supplementary Figure S3.**
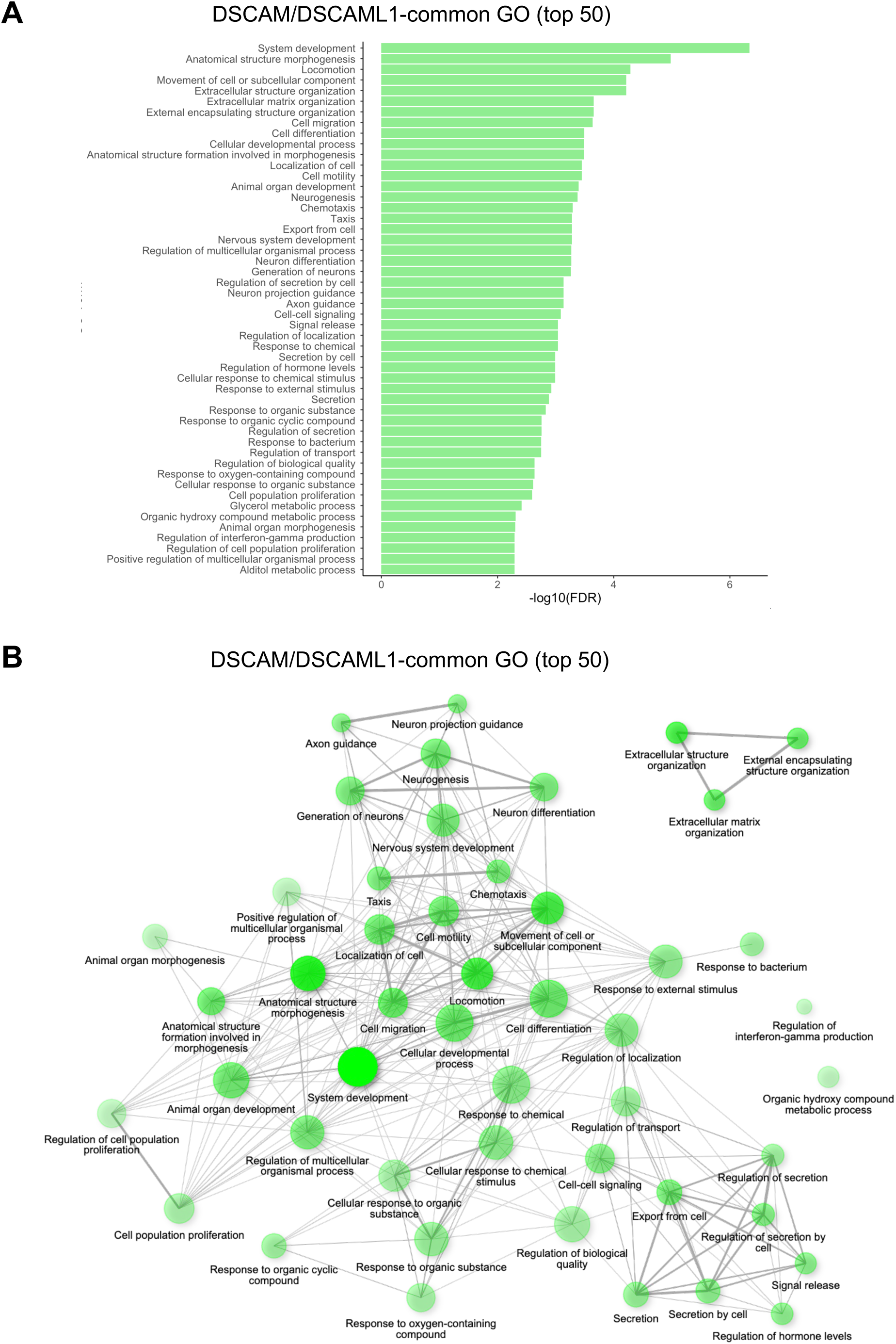
Gene ontology enrichment analysis of differentially expressed genes commonly regulated by overexpression of DSCAM and DSCAML1. (**A**) The top 50 GO enriched items in Biological Process for common DEGs of DSCAM and DSCAML1 performed with ShinyGO. (**B**) The network of the top 50 GO enriched items in Biological Process shown in (**A**). Each node represents an enriched GO term. Related GO terms are connected by a line, whose thickness reflects percent of overlapping genes. The size and the color shade of the node correspond to number of genes and statistical significance, respectively.

**Supplementary Figure S4.**
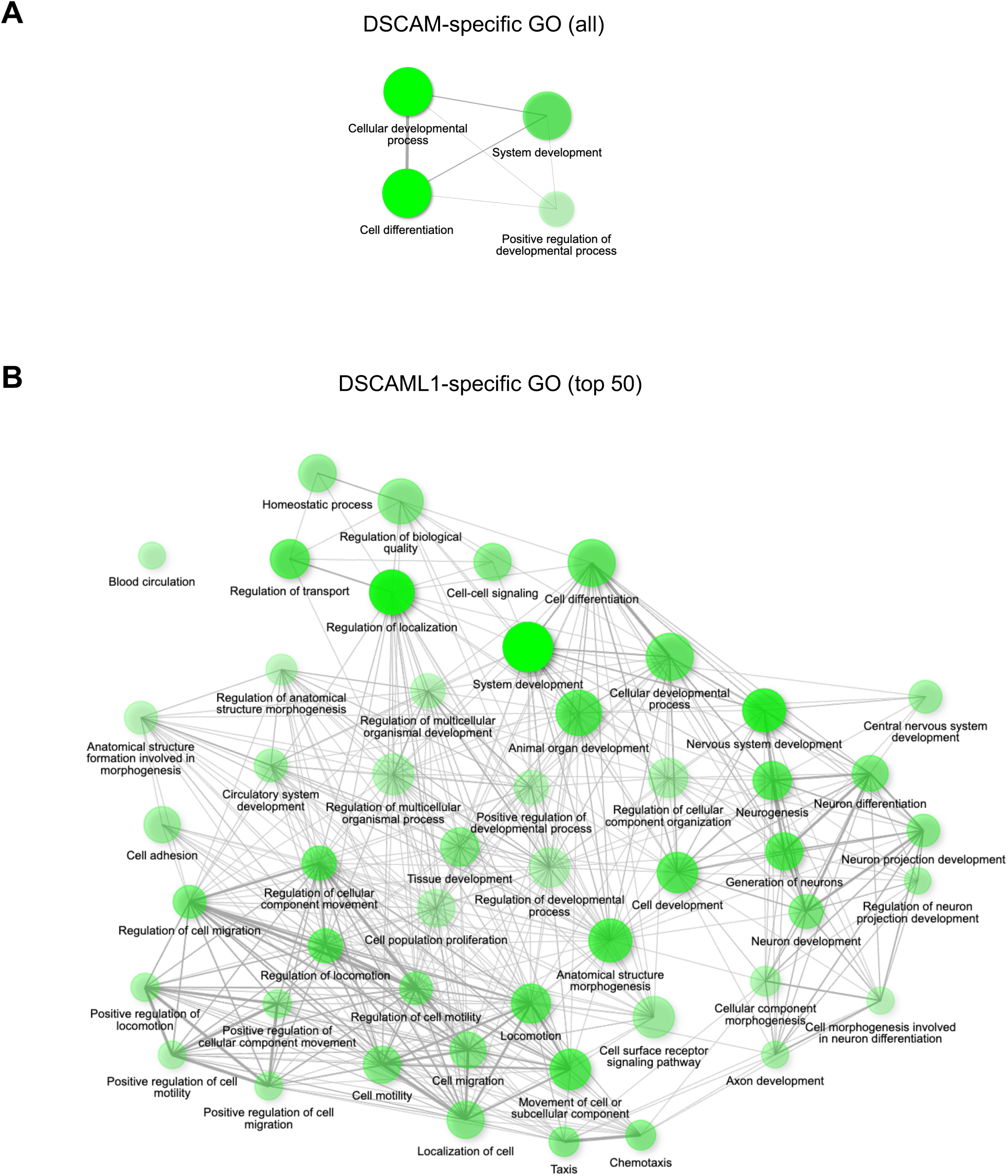
Network visualization of GO biological processes enriched in DSCAM- and DSCAML1-specific differentially expressed genes. (**A**) The network of the all GO terms in DSCAM-specific DEGs. (**B**) The network of the top 50 GO terms in DSCAML1-specific DEGs. Each node represents an enriched GO term. Related GO terms are connected by a line, whose thickness reflects percent of overlapping genes. The size and the color shade of the node correspond to number of genes and statistical significance, respectively.

